# Non-guided, Mobile, CBT-I-based Sleep Training in War-torn Ukraine: A Feasibility Study

**DOI:** 10.1101/2024.08.26.609792

**Authors:** Anton Kurapov, Jens Blechert, Alexandra Hinterberger, Pavlos Topalidis, Manuel Schabus

**Author notes:** Corresponding author: Anton Kurapov.

## Abstract

**Objectives:** To study whether a mobile, unguided Cognitive Behavior Therapy-based Intervention for Sleep Disturbance, Sleep^2^ is feasible, acceptable, and reduces mental health/sleep disturbance symptoms among the Ukrainian population during the ongoing war.

**Methods:** A single-arm, open-label, uncontrolled pre-post evaluation study was conducted with 487 registered participants: 283 started, out of which 95 completed without an ambulatory heart rate (HR) sensor and 65 with. Assessments were conducted using online questionnaires and continuous objective measurements via HR sensors. Key outcome measures included sleep disturbance, insomnia, fear of sleep, anxiety, depression, PTSD, perceived stress, and somatic symptoms.

**Results:** Engagement with the program was robust, achieving an 80.72% compliance rate, alongside high levels of feasibility and acceptance. Participants reported significant pre-post reductions in the severity of sleep disturbance (by 22.60%), insomnia (by 35.08%), fear of sleep (by 32.43%), anxiety (by 27.72%), depression (by 28.67%), PTSD (by 32.41%), somatic symptoms (by 24.52%), and perceived stress (by 17.90%), all with medium to high effect sizes. Objective sleep measurements showed a slight reduction in sleep onset latency.

**Conclusion:** The ‘Sleep^2^Ukraine’ program demonstrated high feasibility and acceptance, with significant improvements in subjective sleep and mental health measures among participants. These findings demonstrate the potential of scalable mobile-based CBT-I interventions in war-torn regions with or without the instrument, based on the heart rate assessment.

## Introduction

### Sleep and Mental Health in Times of War

War can be referred to as a specific form of crisis that inevitably has an impact on various aspects of life (1), including mental health (2) in both military personnel and civilians. Besides typical trauma-related disorders such as anxiety, depression, PTSD, civil victims of war were shown to develop long-term sleeping problems that may last for years even after the conflict termination (3,4). Thus, as for sleeping problems, war has a significant impact on health via (at least) two pathways: through direct causation of trauma related disorders, as well as indirectly by disturbing resilience factors such as healthy and restorative sleep. It is also well known that sleep issues reinforce mental health complaints (5), with unrestorative sleep being the key maintenance factor for anxiety, depression, and PTSD (6). In turn, improvement of sleep quality usually improves mental health symptoms in general (7,8).

As of July 2024, Ukraine is in war for more than two years. Respective recent studies showed significant deterioration of mental health in Ukraine: from the very beginning of the war (9,10) and throughout its progression (11,12). A specific problem seems to be the constant and unpredictable air attacks on the entire territory of Ukraine (13,14). Being mostly carried out during nighttime to complicate air defense, these attacks disturb sleep directly through loud raid alarm systems and likely contribute to fear of sleep through a build-up of anxious apprehension and anticipation of further disturbances and the need to rush for shelter quickly or search for a room without windows at home. Therefore, the need for easily accessible and scalable sleep coaching for Ukrainians is evident and of a high need.

### Mobile-Based Sleep Intervention

One of the most efficient sleep interventions is classical, face-to-face, cognitive behavioral therapy for insomnia (CBT-I) (15). European (16) and US (17) clinical guidelines recommend CBT-I as the first line of treatment. It is a non-pharmacological, well-established intervention, teaching relaxation techniques, sleep restriction, stimulus control, psychoeducation, and cognitive strategies (16). Recent meta-analyses show overall high efficiency of such non-pharmacological, conventional, face-to-face interventions (7,18). Digital CBT-I (dCBT-I) represents the online implementations of these successful programs, with the advantage of being ubiquitously available and cost effective (19). Also, dCBT-I has been studied and has been shown to generate mostly comparable effects to its face-to-face counterpart. In traumatized populations it showed improvements not only on sleep parameters (20,21), but also on other mental health indices (18,22). As such, dCBT-I has also been successfully used in prevention (23).

### Digital CBT for Sleep Disturbances in Ukraine

As indicated above, one of the major advantages of mobile-based CBT interventions in Ukraine is their low threshold. The war conditions make mobility to specialized treatment centers difficult in some areas. Mental health resources are limited and focused on the most acute, physical condition, and the armed forces. Thus, mobile interventions provide access for everyone owning a smartphone with internet connection, thus increasing accessibility, feasibility, and likely acceptance. Yet, due to the poor knowledge of English, treatments offered in Ukrainian language are likely to increase feasibility and acceptance. To our knowledge, ‘Sleep^2^Ukraine’ is the only scalable mobile sleep training delivered to Ukrainians that specifically targets sleep disturbances and comes in Ukrainian language. The novel, evidence-based smartphone application Sleep^2^ (formerly called NUKKUAA®) comprises an unguided, fully automated CBT-I based sleep training program. One particular aspect of the application is the heart rate-based sleep scoring: using chest-belt-derived heart rate data and server-based matching algorithms, sleep stages can be inferred with near to human-expert classification accuracies (24,25). In the morning, objective sleep architecture data are fed back to users, along with norm data comparisons and respective tailored strategies for sleep improvement. Yet, continuous sensor wearing can be bothersome, uncomfortable, and, generally, unguided interventions are known to yield only limited and short-term compliance (26).

### The Present Study

Thus, in preparation for a formal randomized controlled efficacy trial, the present research aimed to assess the uptake, feasibility, acceptance, and usability of the Ukrainian version of Sleep^2^, provided free of charge to all Ukrainian users. The study was preregistered (see the link for details on hypotheses https://osf.io/2gh93): duration had to be extended from 4 to 6 weeks due to a non-daily usage of the program by participants and the necessity to complete most of the offered modules before we conduct post-assessment. Uptake was of a particular relevance as roll out in a population under constant exposure to war can be challenging, particularly in connection with ambulatory biofeedback devices such as the heart rate sensor. Thus, we tested our preregistered hypotheses expecting high levels of uptake, feasibility, and acceptance. We expected self-selection to result in participants with heightened levels of sleep disturbance, our target population, and some mental health symptoms. Further, we predicted improvement of subjective sleep quality and objective sleep parameters such as sleep onset latency, number of awakenings, and sleep efficiency. We also collected some qualitative data on app use. Last, given the interrelations of sleep with mental health and potential generalization of the included CBT techniques, and we also predicted improvement on mental health symptoms (anxiety, depression, post-traumatic stress disorder, somatic symptoms).

## Methods

### Study Design and Procedure

This study was planned as a single-arm, open-label, uncontrolled pre-post evaluation study (see Figure 1). After t0 measures, accommodation and adaptation baseline was followed by t1 measures and the 6 weeks CBT-I program, before t2 measures concluded the study.

**Figure 1.**
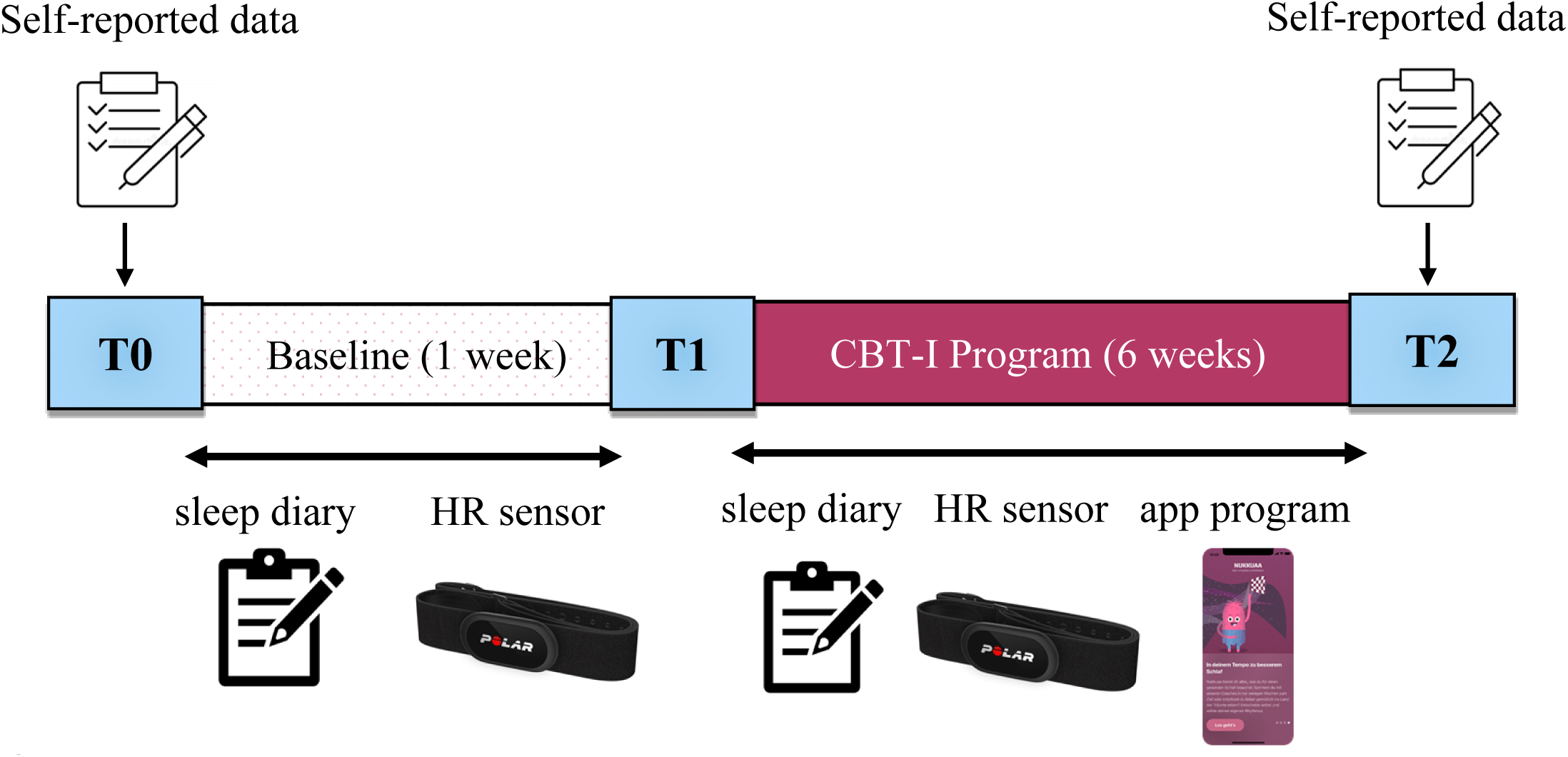
‘Sleep^2^Ukraine’ Study Design. The effects of the CBT-I-based smartphone app (former NUKKUAA®, now sleep²) on sleep were assessed subjectively via questionnaires (at T0 and T2) as well as a sleep diary every morning and objectively via a heart-rate sensor during the study. HR sensor – heart rate sensor.

Daily sleep quality data were assessed continuously between t0 and t2: every morning participants had to fill in the sleep diary. Due to the fully anonymous mode of data collection we could not prompt or encourage participants throughout training in case of missing diaries. Also, after training discontinuation at t2, we lost contact and could not conduct follow-up data, neither could we follow-up on dropouts. Recruitment and registration for the study started on October 1^st^, 2023 and ended on November 1^st^, 2023. During registration, participants could voluntarily choose whether they want to receive the heart rate (HR) sensor free of charge, but with the necessity to return it after 6 weeks at the latest. To maintain anonymity, the sensor was shipped to participant through a third-party service provider^1^. All participants have provided written informed consent at t0, which is documented in the survey.

### CBT-I-based App Program

The Sleep^2^ app content was translated, rewritten to fit Ukrainian language grammar, and culturally adapted (27). The program is organized into 6 consecutive levels, each featuring a set of components, based on core elements of CBT-I. It features various modes of communication, e.g., videos teaching sleep hygiene and psychoeducation, a chat-bot designed to help reframe thoughts, audio exercises for relaxation, tips on keeping good sleep habits, and blog posts discussing important scientific sleep topics (28). For details of the program content, see suppl.Table 1. During the 6-week app phase, participants were instructed to complete the program’s six levels, which are designed to contain the most important contents of CBT-I. At t2 they were redirected to post-treatment online questionnaires directly from the app. Participants not reaching level 6 and completing t2 measures were considered dropouts (we did not obtain information on partial completion).

**Table 1.** Sociodemographic Information, Participant Flow, Engagement, and Compliance.

| Sociodemographics |  |  |  |  | Engagement (%) | Compliance (%) |  |  |
| --- | --- | --- | --- | --- | --- | --- | --- | --- |
| Participants | N | Female sex (%) | Age (M ± SD) | Financial Status (€) | Moring Diaries | Exercises Completed | Educational Videos Watched | Chat-bot Completed |
| Registered | 487 | 71 |  |  |  |  |  |  |
| Non-starters | 204 | 73 |  |  |  |  |  |  |
| Starters | 283 | 77 | 27.9 ± 9.15 | 115.7 ± 89.2 | 35.68 | 51.27 | 8.89 | 8.21 |
| Non-completers | 123 | 77.7 | 29.2 ± 10.27 | 116.4 ± 112.1 | 30.41 | 37.69 | 7.26 | 6.13 |
| Completers | 160 | 76.9 | 27.3 ± 8.14 | 123.02 ± 88.6 | 38.49 | 57.25 | 9.78 | 8.96 |
| No Sensor | 95 | 82.5 | 27.7 ± 9.51 | 161.9 ± 130.8 | 35.16 | 57.34 | 10.65 | 9.94 |
| With Sensor | 65 | 64.6 | 28.1 ± 6.72 | 126.7 ± 99.9 | 39.63 | 54.53 | 7.88 | 7.51 |

*Note.* The table represents average values per every group of participants. Compliance is calculated based on the total number of entries from a single participant of daily audio exercises completed (had to be performed minimum one time per day), psychoeducational videos and chat (had to be performed at least once per level).

## Material and Measures

### Acceptance, Engagement, and Compliance

**Acceptance** was measured with an adapted 12-item self-report measure about participants’ experiences and impressions of the program (suppl.Table 3). Participants gave information on app usability, program helpfulness, content relevance, helpfulness of the push-notifications, frequency of the reminders, and program duration using a 10-point Likert-type scale ranging from 1 (strongly disagree) to 10 (strongly agree).

**Engagement** was monitored through the app’s log data (frequency and duration of app use). Participants were encouraged to engage with the app via daily push-notifications.

**Compliance** was measured by the adherence to the program’s requirements: participants were required to complete at least five audio exercises, one video, and one chatbot interaction per each of the 6 levels. We also measured the sleep diary entries.

### Self-report Measures

Self-reported data was obtained with online questionnaires at t0 and t2 focusing on a time range of one month prior to the date of filling out the questionnaire. Participants were required to respond to all questions, thus yielding no missing values. If no Ukrainian adapted version of the questionnaire was available, items were translated by the authors. For details on scoring ranges, number of items, and reliability statistics see suppl.Table 2.

**Table 2.** Summary of the Screened Symptoms at Baseline (t0) for Completers (N=160)

| Symptom | Mean | SD | Cutoff | Count | Percentage |
| --- | --- | --- | --- | --- | --- |
| Sleep Disturbance | 8.93 | 3.45 | No sleep issues | 16 | 9.69 |
|  |  |  | Sleep disturbance | 99 | 60 |
|  |  |  | Chronic sleep problems | 50 | 30.31 |
| Insomnia | 11.98 | 5.65 | Subthreshold insomnia | 74 | 44.85 |
|  |  |  | Clinical insomnia of moderate severity | 46 | 27.88 |
|  |  |  | Severe clinical insomnia | 7 | 4.24 |
| Fear of Sleep | 5.55 | 6.67 | Mild fear | 21 | 12.73 |
|  |  |  | Moderate fear | 8 | 4.85 |
|  |  |  | Severe fear | 1 | 0.61 |
| Somatic Symptoms | 9.42 | 4.73 | Minimal symptoms | 15 | 9.09 |
|  |  |  | Low symptoms | 45 | 27.27 |
|  |  |  | Medium symptoms | 55 | 33.33 |
|  |  |  | Severe symptoms | 31 | 18.79 |
| Anxiety | 7.36 | 3.81 | Mild anxiety | 76 | 46.06 |
|  |  |  | Moderate anxiety | 43 | 26.06 |
|  |  |  | Severe anxiety | 0 | 0 |
| Depression | 11.82 | 5.48 | Mild depression | 48 | 29.09 |
|  |  |  | Moderate depression | 52 | 31.51 |
|  |  |  | Moderately severe depression | 33 | 20 |
|  |  |  | Severe depression | 19 | 11.51 |
| PTSD | 27.27 | 15.21 | Presence of PTSD | 10 | 6.06 |
|  |  |  | Severe PTSD | 47 | 28.48 |
| Perceived Stress | 7.38 | 3.18 | - | 95 | 57.57 |

*Note.* The table presents mean values and standard deviations for all measured symptoms, together with the clinical cutoffs per every measure, number of participants and corresponding percentage.

Sociodemographic data were obtained on sex, age, employment status, working conditions, financial situation, and internally displaced persons (IDP) status.

### Sleep-Related Questionnaires

Sleep disturbance was measured with the Ukrainian version of the 19-item Pittsburgh Sleep Quality Index (PSQI, Mazur et al., 2021). Insomnia symptoms were measured with the 7-item Insomnia Severity Index (ISI; Bastien, 2001). Fear of Sleep in the past month was measured with the 13-item Fear of Sleep Inventory (FoSI; Pruiksma et al., 2014).

### Mental Health Symptom Questionnaires

**Anxiety** was measured with the Ukrainian version of the 7-item General Anxiety Disorder-7 scale (GAD-7; Shyroka & Mykolaychuk, 2020). **Depression** was measured with the Ukrainian version of Patient Health Questionnaire- 9 (PHQ-9; Unifikovanyy Klinichnyy Protokol Pervynnoyi, Vtorinnoyi (Spetsializovanoyi) Ta Tretynnoyi (Vysochospetsializovanoyi) Medychnoyi Dopomohy, 2014). **PTSD** symptoms were measured with the 20-item Ukrainian version of The PTSD Checklist for DSM-5 (PCL- 5; Karachevskii, 2016). **Perceived Stress** was measured with the 4-item Perceived Stress Scale (PSS-4; Warttig et al., 2013). **Somatic Symptoms** were assessed with the 8-item Somatic Symptom Scale (SSS-8; Gierk et al., 2014).

### Objective Sleep Measurement

In participants receiving and using the HR sensor Polar® H10 (Polar Electro GmbH Deutschland), continuous objective sleep measures were available. HR data were read out in the morning in combination with the subjective morning sleep diary, transferred to the server via https protocol and processed using the deep learning network as described in Topalidis, Baron, et al. (2023) and Topalidis, Heib, et al. (2023), resulting in a detailed sleep analysis. Continuous sleep measurements included time in bed (TIB), total sleep time (TST), sleep onset latency (SOL), number of awakenings (NOA), wake after sleep onset (WASO), and sleep efficiency (SE).

### Statistical Analysis

Daily/nightly data were averaged over the baseline (first 7 days of the study between t0 and t1) and over the last 7 days before the end of the treatment (t2, see Figure 1) for statistical analysis, as performed in R, version 2023.03.0 (37). Extreme outliers, that is, data points deviating more than ±2 *SD* from the mean, were excluded from the respective analysis. Besides completer analyses we also report intention-to-treat (ITT) analyses (with pre-assessment values (t1) were carried forward to the post-assessment (t2)).

## Results

### Recruitment Flow and Program Uptake

A total of 487 people initially registered for the sleep training, out of which 283 started (t0) the program, and 160 finished (t2): 95 without continuous objective measurements and 65 with the sensor (see Table 1) resulting in a completion rate (t0-t2) of 56.54%. As can be seen in Table 1, there were no obvious differences between starters and completers on sociodemographics, but completers evidently completed more content. The analysis of the dropouts (N = 123) in comparison to program completers (N = 160) revealed no difference in age (*t*(250) = 0.45, *p* = .651), sex (*χ^2^*(3) = 2.96, *p* = .398), employment status (*χ^2^*(1) = 0.25, *p* = .619), working conditions (*χ^2^*(2) = 0.37, *p* = .829), displacement status (*χ^2^*(3) = 2.52, *p* = .472), and financial situation (*χ^2^*(7) = 8.78, *p* = .269). Interestingly, those participants demanding and receiving the HR sensor were more likely to be male and had less favourable financial situation. Yet, critically, they were not consistently more engaged with the program.

At t0, completers reported high levels of sleep disturbance, insomnia, fear of sleep, anxiety, depression, PTSD, perceived stress, and somatic symptoms (see Table 2), confirming that we attracted our target population. On average, 6.36 (*SD* = 13.51) out of 40 nights were disturbed by air-raid alarms.

### Engagement: Session Completion and Overall App Use

Overall, the N=160 completers saved 80.72% of the 50 required sleep diaries, representing good compliance. Completers used the app on an average of 39.8 days (*SD* = 19.12) out of a maximum of 50 days.

### Usability and Program Acceptance

Most of the participants were very satisfied with the treatment and the app, specifically content clarity, content relevance, usefulness of the relaxation exercises, relevance of the proposed program levels, and with recommending the program to others (see Table 3).

**Table 3.** Summary of the Program Evaluation.

Summary of the Program Evaluation
| Category | Mean | SD |
| --- | --- | --- |
| Content clarity | 8.69 | 1.88 |
| Recommend Sleep<sup>2</sup>Ukraine to others | 8.17 | 2.07 |
| Content relevancy | 7.64 | 2.2 |
| Usefulness of relaxation exercises | 7.6 | 2.68 |
| Levels relevancy | 7.22 | 2.52 |
| Convenience to sleep | 7.03 | 2.47 |
| Improved sleep | 6.49 | 2.27 |
| Usefulness of imaginary exercises | 6.69 | 2.99 |
| Sleepscore accuracy | 6.67 | 2.43 |
| Easy to meet requirements for next level | 6.38 | 2.84 |
| Personalized program effectiveness | 6.25 | 2.29 |
| Usefulness of push-notifications | 5.8 | 3.25 |
| Usefulness of the chatbot | 5.38 | 3.14 |
| Problems using sensor | 4.99 | 3.13 |
| Wake up because of sensor | 3.01 | 2.86 |

*Note.* The table presents the average survey results on the acceptance of the proposed program, sorted in descending order. Participants rated the app and the sensor on specified criteria using a 10-point Likert scale (1 = strongly disagree to 10 = strongly agree). The evaluation intervals are as follows: very bad (0-2), bad (3-4), mediocre (5-6), good (7-8), very good (9-10). Formulations of items is presented in Suppl.Table 3. Results are presented for the sensor users (N = 65).

As for **qualitative feedback** participants reported that the sensor restricted sleeping positions, was uncomfortable to adjust and wear, and had occasional synchronization and battery issues. While many found the objective sleep analysis helpful (78%), others doubted its accuracy (12%). Many (56%) participants preferred a wrist/arm sensor instead of a chest belt. Positive feedback highlighted ease of use, diverse content, helpful relaxation exercises (especially before sleep and after air raid alarms), pleasant design, simple interface, engaging educational content and chatbot, and the ability to take and reflect on notes within the app.

Negative feedback included limited exercise variety, inability to revisit or list all exercises, lack of video adjustments (speed, narrator, subtitles), no late sleep entries or nap recordings, limited chatbot responses, and a discouraging level structure suggesting continuous sleep improvement; a topic-based division was preferred.

### Pre-post Changes on Sleep and Mental Health

Completers reported statistically significant improvements on all symptoms after the sleep training including sleep disturbance, insomnia, fear of sleep, anxiety, depression, PTSD, perceived stress, somatic symptoms, and resilience with medium to high effect sizes (Table 4, see also density distributions and clinical cutoffs in Supl.Figure 1). Similar results were found in the ITT analysis (N = 283; see Supl. Table 4), which speaks against selective dropout effects.

**Table 4.** Changes in Mental Health Symptoms after the Sleep Training (N=160 completers)

|  | **T0** | **T2** |  |  |  |  |
| --- | --- | --- | --- | --- | --- | --- |
|  | **M ± SD** | **M ± SD** | **T2-T0** | **t(159)** | **p** | **Cohen's d** |
| Sleep Disturbance | 8.93 ± 3.45 | 6.91 ± 3.02 | -2.02 | 6.64 | <.001 | .53 |
| Insomnia | 11.98 ± 5.65 | 7.78 ± 4.84 | -4.20 | 8.75 | <.001 | .69 |
| Fear of Sleep | 5.55 ± 6.67 | 3.75 ± 5.83 | -1.80 | 3.12 | .002 | .25 |
| Anxiety | 7.36 ± 3.81 | 5.34 ± 3.21 | -2.02 | 6.05 | <.001 | .48 |
| Depression | 11.82 ± 5.48 | 8.43 ± 4.74 | -3.39 | 6.52 | <.001 | .52 |
| PTSD | 27.27 ± 15.21 | 18.42 ± 13.97 | -8.85 | 6.34 | <.001 | .51 |
| Stress | 7.38 ± 3.18 | 6.06 ± 2.74 | -1.32 | 4.92 | <.001 | .39 |
| Somatic Symptoms | 9.42 ± 4.73 | 7.10 ± 4.19 | -2.32 | 6.34 | <.001 | .51 |

*Note.* The table presents the results of paired-sample t-test for subjective measures obtained with the questionnaires at T0 and T2 (see Figure 1 for details). Results are presented for program completers (N = 160).

### Objective Sleep Measures (Sensor Users Only)

Table 5 shows that the N=65 program completers with sensors showed small improvements on objective SOL (reduction of time to fall asleep from ∼29 to 25 min).

**Table 5.** Changes in Continuous Sleep Measures from pre (t1) to post (t2) Training.

|  | **T1** | **T2** |  |  |  |  |
| --- | --- | --- | --- | --- | --- | --- |
|  | **M ± SD** | **M ± SD** | **T2-T1** | **t(73)** | **p** | **Cohen's d** |
| TIB | 492.45 ± 90.37 | 489.69 ± 82.77 | 4.84 | 0.891 | .376 | .11 |
| TST | 427.17 ± 77.95 | 428.74 ± 80.26 | -0.91 | -0.180 | .858 | -.02 |
| SOL | 29.71 ± 33.34 | 25.01 ± 29.28 | 4.71 | 2.047 | .044 | .24 |
| NOA | 2.44 ± 2.45 | 2.42 ± 2.35 | 0.06 | 0.403 | .688 | .05 |
| WASO | 40.49 ± 33.69 | 40.56 ± 36.84 | 0.66 | 0.205 | .838 | .02 |
| SE | 86.93 ± 8.03 | 87.66 ± 8.63 | -0.83 | -1.368 | .176 | -.16 |

*Note.* NOA (number of awakenings) here is a count of awakenings of more than 2 continuous minutes. TIB – time in bed, TST – total sleep time, SOL – sleep onset latency, WASO – wake after sleep onset, SE – sleep efficiency. Results are presented for sensor users (N = 65).

## Discussion

To our knowledge, this is the first study to report on a non-pharmacological, digital CBT-I intervention during an ongoing war. The results, obtained from Ukraine, demonstrated high uptake/feasibility and acceptance of the program, with significant improvements across all sleep and mental health measures. We will discuss each of these in this order in the following.

### Uptake, Feasibility, Acceptance, Usability

*Program uptake* was moderate: out of 487 registered users, 283 started. Out of 283 starters 160 completed, thus, a completion rate was 56%, which can be considered very good, given the circumstances and the high demands of the program (6 levels, 50 required diaries). This means that recruitment for the subsequent randomized controlled trials (RCT) should be very feasible. Interestingly, despite additional logistics via mail for the sensor and the discomfort reported by some, 65 out of the 160 program completers proceeded with the daily use of the HR sensor.

Regarding *engagement*, participants used the app for a duration of 6 weeks, with 57% completing the full training and showing above-average compliance and engagement. The overall dropout rate of 43.46% is similar to other studies investigating the effects of digital programs for insomnia in outpatients: 40% (38) and 34.4% (39). In fact, most participants completed at least half of the program (∼20 days), with no obvious demographic differences between dropped-out participants and program completers.

Whereas veridical feedback on objective sleep parameter can be very helpful for insomnia, the delivery of a HR sensor is an obvious hurdle to uptake and to largescale roll- out. Given our non-randomised assignment to sensor vs. non-sensor groups we can only hint at these effects. Sensor users were more likely to be male and had less favourable financial situation. The further might reflect some gender-stereotypical technology affinity in Ukraine users, whereas the latter points to the potentially higher subjective value of receiving a costly wearable. Importantly, we cannot conclude that the sensor users were more engaged. Should this be confirmed in further studies with random assignment to sensor vs. no-sensor groups then we could remove this hurdle to large scale application (at the cost of objective sleep data).

### User satisfaction and potential app improvements

Those who completed the training and moved to t2 questionnaires were highly satisfied, especially with the (culturally adapted) training content, relaxation exercises, assigned levels, and were willing to recommend the program. The lowest scores were for the chatbot and push notifications, indicating these features were less effective. As for qualitative feedback, only some sensor users reported significant discomfort. Objective feedback was considered useful, with positive remarks on relaxation exercises and negative feedback on their limited variation and inability to revisit previous exercises. Participants also disliked the inability to record daytime naps and/or sleep shorter than four hours. However, this is a technical limitation as the algorithm has not been trained on nap data and, therefore, can at that point not be verified.

The positive reception of the intervention and the relatively low dropout rate suggest that there is a significant demand and openness to internet-based interventions among people in Ukraine. This contrasts with the situation in Western countries, where internet interventions often face challenges with low uptake (40,41). One possible reason for this difference is the low availability of mobile interventions in the Ukrainian language. This indicates a general reluctance among Ukrainians to use interventions available in English or other languages.

This insight has important implications for the development of interventions in other mental health areas for Ukraine: language accessibility appears to be a crucial factor.

### Sample Characteristics, Pre-Post Effects on Sleep Parameters and Mental Health

#### Participant Flow and Symptom Severity

Regarding the pathology of the sample and whether we addressed the program’s target group: our sample demonstrated significant sleep impairments, confirming that we reached the intended population. Before starting the training, participants who completed the program exhibited significantly elevated levels of sleep disturbance (60% of the sample), with 30% experiencing chronic sleep problems, subthreshold and clinical insomnia (44 and 27%, respectively), figures that are nearly three times higher than those normally observed in Western populations (28,42). We also observed very high levels of severe PTSD among participants, potentially reflecting the ongoing air war on civilians, which highlights the necessity for PTSD-specific content in the program. In addition to this, more than 52 percent of our participants had elevated somatic symptoms, 60 percent mild to moderate anxiety or depression, and more than half reporting elevated levels of stress.

### Pre-Post Effects on Mental Health Symptoms and Sleep Parameters

Program completers reported substantial reductions in subjective sleep disturbance (minus 22.60%), insomnia severity (minus 35.08%), and fear of sleep (minus 32.43%), all with medium to high effect sizes. There were also notable decreases in anxiety (minus 27.72%), depression (minus 28.67%), PTSD (minus 32.41%), somatic symptoms (minus 24.52%), and perceived stress (minus 17.90%). In sensor-wearing completers, objective sleep measurements did not show large scale significant changes post-training, except for a slight reduction in objective sleep onset latency (SOL). Given that the baseline objective sleep data indicated no significant sleep disturbances (with total sleep time above 7 hours), the lack of objective changes over the 6-week study period is not surprising. The striking discrepancy between subjective and objective sleep measures is a known issue, particularly in insomnia (43). Our findings reiterate the predominantly subjective nature of many sleep disorders (44).

Our results align with previous findings (45), highlighting the strong co-occurrence between sleep disturbance and somatic symptoms (46), depression and anxiety (7,47), and PTSD (48). A main finding of our study shows that all trauma-related disorders improve over the course of a predominantly sleep-focused program. This might be related to the above-mentioned indirect pathways: as sleep recovers, also depression and PTSD might do so. Yet, many of the taught CBT-based cognitive techniques are applicable to these non-sleep related symptoms just as well. Further, self-efficacy related to sleep improvements might translate onto other domains as well.

### Limitations and Future Directions

Our study has several *limitations*. First, it used a convenience sample and lacked a control group, being a single-arm, open-label, uncontrolled pre-post evaluation study. More comprehensive follow up studies using randomized group assignment and including control groups are necessary to rule out that improvements are not happening also in the absence of any intervention, simply due to passing time, measurement repetition, symptom awareness or any other factors. However, in a war-torn population, it is ethically challenging to use an inactive control or waitlist group, which, together with the fully anonymous data, also led to the absence of the follow-up. Secondly, the lack of representative random sampling may introduce selection bias, as participants who self-subscribed to the sleep training may have more severe sleep problems and/or mental health issues and a higher affinity for digital interventions than the general population.

Scalable mobile-based interventions seem crucial for the Ukrainian population, especially in war-threatened areas with available internet and electricity. Our study demonstrates that such interventions can be effectively delivered. Participants are largely reachable and ready to accept help. CBT-I is simple, safe, and effective, showing promising effects within 6 weeks. Given the high feasibility and acceptability, further research, including randomized controlled trials, is necessary to verify the efficiency of mobile-based CBT-I in Ukraine.

## Author contribution

AK – study design, data collection, data analysis, manuscript writing JB – study design and methodology, manuscript review and editing AH – manuscript review and editing

PT – manuscript review and editing

MS – study design and methodology, manuscript review and editing

## Acknowledgments

Research was funded by MSCA4Ukraine, a funding scheme that is implemented by a consortium comprised of Scholars at Risk Europe (SAR Europe) hosted at Maynooth University, Ireland (project coordinator), the German Alexander von Humboldt Foundation (AvH) and the European University Association (EUA); grant number 1233157.

IBI-based algorithms for sleep classification were kindly provided for scientific purposes by sleep² (Nukkuaa GmbH).

## Data Availability Statement

The data for analysis of this study are provided on OSF repository (https://osf.io/2gh93).

## Ethics Statement

The studies involving human participants were reviewed and approved by the Ethics Committees of Paris Lodron University of Salzburg (EK-GZ 26/2023) and Taras Shevchenko National University of Kyiv (11-22/7). The participants provided their written informed consent to participate in this study.

## Supplemental Material

**Supplementary Figure 1.**
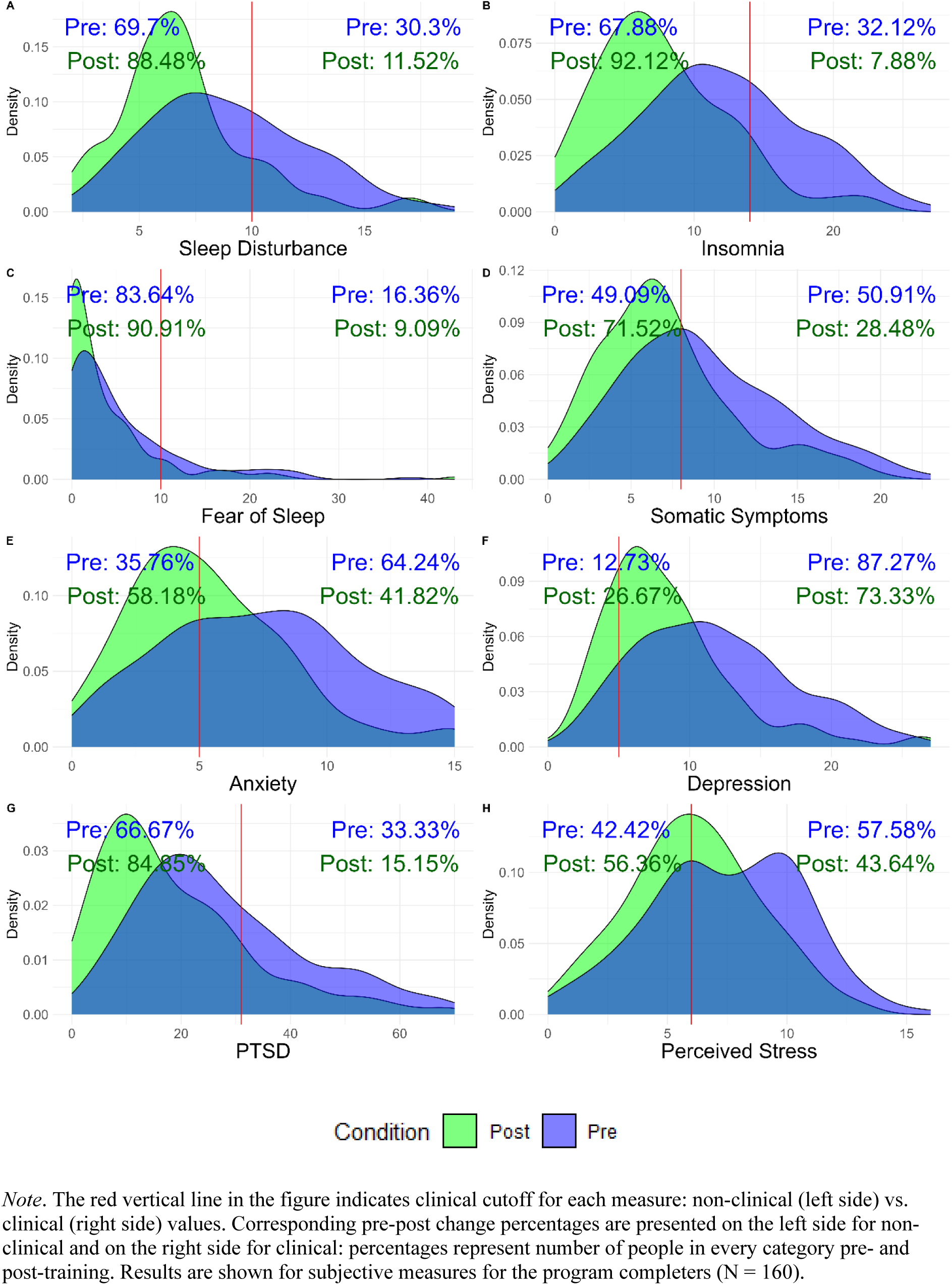
Density Differences between Pre- and Post-Measurements for Program Completers (N = 160)

**Supplementary Table 1.** Overview over the Contents of the App Program Divided by Level (28)

| **Level** | **Video on sleep education** | **Auditory relaxation exercise** | **Sleep training via chat-bot** | **Blog** | **Sleep hygiene advice** |
| --- | --- | --- | --- | --- | --- |
| **1** | Effects of tension and relaxation on sleep | Progressive muscle relaxation (PMR) | Topic: Rumination + exercise to putting those thoughts aside | 1. Correlations between sleep and depression/anxiety<br>2. Sleep and (cognitive) performance | „Go to bed only when you are tired” |
| **2** | Education on healthy sleep + sleep cycles | Imagination exercise | Topic: What to do with the “new free time” that was previously spent awake in bed | 1. Why do we need healthy sleep<br>2. Sleep, mental health, and online therapy | „After lying awake for 20 minutes, get up and return to your bed once you feel tired again” |
| **3** | Circadian rhythms | Breathing exercise | Topic: Sleep restriction | 1. Powering through the week and sleeping in at the weekend - a good idea?<br>2. Chronotype | „Only spend as much time in your bed as you’ll actually sleep” |
| **4** | Factors influencing and disturbing healthy sleep | Imagination exercise | Topic: Habits influencing good sleep | 1. Insufficient sleep and impacts on health<br>2. Negative effects of electromagnetic radiation on sleep | „The bed should only be for sleeping (and sleeping with other people)” |
| **5** | Sleep and medication | Imagination exercise | Topic: Mindfulness | 1. Sleep medication<br>2. Natural sleep aids | „Try to have a regular sleep-wake-rhythm” |
| **6** | Sleep throughout the course of life | Meditation | Topic: Bedtime routines | 1. Age and sleep<br>2. Changes in sleep-wake-rhythms throughout the course of life | „Spend 30 minutes outside every day” |

**Supplementary Table 2.** Scoring Ranges, Number of Items, and Reliability Statistics for Psychological Questionnaire Measures.

| Variable Group | Variable | Measure | Reliability (Cronbach's Alpha) |  | Range (min-max) | Number of Items |
| --- | --- | --- | --- | --- | --- | --- |
|  |  |  | T1 | T2 |  |  |
| Sleep | Sleep Disturbance | PSQI | .72 | .74 | 0-21 | 19 |
|  | Insomnia | ISI | .84 | .83 | 0-28 | 7 |
|  | Fear of Sleep | FoSI | .84 | .86 | 0-65 | 13 |
| Mental Health | Anxiety | GAD-7 | .88 | .89 | 0-21 | 7 |
|  | Depression | PHQ-9 | .84 | .83 | 0-27 | 9 |
|  | PTSD | PCL-5 | .93 | .94 | 0-80 | 20 |
|  | Perceived Stress | PSS-4 | .93 | .94 | 0-16 | 4 |
|  | Somatic Symptoms | SSS-8 | .74 | .74 | 0-32 | 8 |
|  | Resilience | BRS-6 | .72 | .68 | 6-30 | 6 |

*Note.* This table shows names of the variables, corresponding measures, reliability statistics, scoring range, and the number of items per measure.

**Supplementary Table 3.** Formulations of Items Regarding the Acceptance of the Program.

| No. | Question |
| --- | --- |
| 1 | Has the quality of your sleep improved after following the proposed program? |
| 2 | How effective was the proposed program for you personally? |
| 3 | Was the content of the application generally clear to you? |
| 4 | Did the content of the exercises address the problems they were supposed to solve? |
| 5 | Did the "sleep quality index" calculated in the application match your subjective feelings about the quality of your sleep? |
| 6 | How accurate do you consider the program's level distribution? |
| 7 | Was it easy for you to meet all the level requirements to move to the next one? |
| 8 | Were the relaxation exercises useful for you? |
| 9 | Were the visualization exercises useful for you? |
| 10 | Was the chatbot in the application useful for you? |
| 11 | Were the push notifications (background reminders) useful for you? |
| 12 | Would you recommend 'Sleep<sup>2</sup>Ukraine' to your family, friends, and acquaintances? |

**Supplementary Table 4.** Changes in Symptoms after the Sleep Coaching for All Participants.

|  | **T0** | **T2** | **T2-T0** | **t(281)** | **p** | **Cohen's d** |
| --- | --- | --- | --- | --- | --- | --- |
|  | **(M ± SD)** | **M ± SD)** |  |  |  |  |
| Sleep Disturbance | 8.87 ± 3.58 | 7.67 ± 3.34 | -1.2 | 5.7 | <.001 | 0.34 |
| Insomnia | 11.98 ± 5.69 | 9.40 ± 5.64 | -2.58 | 6.72 | <.001 | 0.4 |
| Fear of Sleep | 5.55 ± 6.75 | 4.12 ± 6.26 | -1.43 | 3.59 | <.001 | 0.214 |
| Anxiety | 7.18 ± 3.71 | 6.03 ± 3.44 | -1.15 | 4.9 | <.001 | 0.292 |
| Depression | 11.76 ± 5.55 | 9.74 ± 5.30 | -2.02 | 5.63 | <.001 | 0.335 |
| PTSD | 27.08 ± 15.42 | 21.73 ± 15.06 | -5.35 | 5.64 | <.001 | 0.336 |
| Stress | 7.46 ± 2.97 | 6.58 ± 2.80 | -0.88 | 4.56 | <.001 | 0.272 |
| Somatic Symptoms | 9.29 ± 4.62 | 7.73 ± 4.16 | -1.56 | 5.1 | <.001 | 0.304 |

*Note.* The table presents the results of paired-sample t-tests for subjective measures obtained with the questionnaires at T0 and T2 (see Figure 1 for details) for all participants (N = 283), including dropouts (N = 123). For dropouts, pre-assessment values were carried forward to the post-assessment, assuming no change due to the intervention, as part of an intention-to-treat analysis in this non-controlled pilot study.

## Footnotes

1 The sensors were shipped to a responsible person in Ukraine, who then contacted participants to gather the necessary details for delivery. The delivery was carried out using the courier service "Nova Poshta" (Eng. "New Post"). Participants were given the option to either collect the belts at the post office or have them delivered directly to their address. As such, we did not know the identities of our participants and could not match their data with their personalities, as study emails were used to log into the app, which did not disclose any personal details, ensuring complete anonymity.

